# Reversible brain edema in experimental cerebral malaria is associated with transcellular blood-brain barrier disruption and delayed microhemorrhages

**DOI:** 10.1101/2021.11.26.470078

**Authors:** Jessica Jin, Mame Aida Ba, Chi Ho Wai, Sanjib Mohanty, Praveen K. Sahu, Rajyabardham Pattnaik, Lukas Pirpamer, Manuel Fischer, Sabine Heiland, Michael Lanzer, Friedrich Frischknecht, Ann-Kristin Mueller, Johannes Pfeil, Megharay Majhi, Marek Cyrklaff, Samuel C. Wassmer, Martin Bendszus, Angelika Hoffmann

**Affiliations:** Department of Neuroradiology, Heidelberg University Hospital, Germany; Division of Experimental Radiology, Department of Neuroradiology, Heidelberg University Hospital, Germany; Centre for Infectious Diseases, Parasitology Unit, Heidelberg University Hospital, Heidelberg, Germany; Center for the Study of Complex Malaria in India, Ispat General Hospital, Rourkela, Odisha, India; Department of Intensive Care, Ispat General Hospital, Rourkela, Odisha, India; Department of Neurology, Division of Neurogeriatrics, Medical University of Graz, Graz, Austria; German Center for Infection Research (DZIF), Heidelberg, Germany; Center for Childhood and Adolescent Medicine, General Pediatrics, University Hospital, Heidelberg, Germany; Department of Radiology, Ispat General Hospital, Rourkela, Odisha, India; Department of Infection Biology, London School of Hygiene & Tropical Medicine, London, United Kingdom; University Institute of Diagnostic and Interventional Neuroradiology, University Hospital Bern, Inselspital, University of Bern, Switzerland

## Abstract

Brain swelling occurs in cerebral malaria (CM) and may either reverse or result in fatal outcome. It is currently unknown how brain swelling in CM reverses, as investigations have been hampered by inadequate animal models. In this study, we show that reversible brain swelling in experimental murine cerebral malaria (ECM) can be induced reliably after single vaccination with radiation-attenuated sporozoites as revealed by *in vivo* high-field (9.4T) magnetic resonance imaging. Our results provide evidence that parenchymal fluid increase and consecutive brain swelling results from transcellular blood-brain barrier disruption (BBBD), as revealed by electron microscopy. This mechanism enables reversal of brain swelling but does not prevent persistent focal brain damage, evidenced by microhemorrhages, in areas of most severe BBBD. In a cohort of 27 pediatric and adult CM patients (n=4 fatal, n=23 non-fatal) two out of four fatal CM patients (50%) and 8 out of 23 non-fatal CM patients (35%) showed microhemorrhages on MRI at clinical field strength of 1.5T, emphasizing the translational potential of the experimental model. Our data suggest that targeting transcellular BBBD may represent a promising adjunct therapeutic approach in cerebral brain swelling to reduce edema and may ultimately lead to a reduced permanent brain damage and a better longtime neurological outcome.

**Author summary:** Brain swelling, which occurs in diseases such as cerebral malaria, is not necessarily fatal, and may reverse. Even upon reversal of brain swelling, neurological sequelae can still occur. The factors contributing to the reversibility of brain edema are not known, and treatment options remain therefore limited. Identifying the mechanisms leading to such reversibility could inform clinical management aimed at decreasing brain swelling and consecutive brain injury. Here we introduce a reproducible and simple animal model that allows comprehensive *in vivo* studies of reversible brain swelling in cerebral malaria at the peak of disease and upon recovery. We identify a specific type of blood-brain barrier disruption (BBBD) as a mechanism that occurs in brain swelling. We show that BBBD can reverse, but also highlight remaining brain damage in areas of most severe BBBD. As the ECM model introduced here bares crucial similarities to the CM in humans, our findings open strategies to study new therapeutic avenues and point to compounds that specifically target transcellular BBBD to reduce brain edema, and increase survival rates.

## Introduction

Cerebral malaria (CM) is the most severe complication of *Plasmodium falciparum* infection. It is characterized by altered consciousness and coma, and by brain swelling in children and, to a lesser degree, in adults. Mortality is high, despite adequate anti-parasitic treatment, and long-term neurocognitive impairment are reported in about one third of surviving patients [1-3].

A hallmark of CM is the sequestration of parasite-infected erythrocytes in the microvasculature, a process described in both human and experimental CM [1, 4, 5]. Pathological mechanisms in CM have been related to endothelial dysfunction and increased blood-brain barrier disruption (BBBD) and it is well accepted that this event precedes vasogenic brain edema [6, 7]. BBBD has long been considered to be caused by disrupted tight junctions (TJs) between endothelial cells in vessels, a phenomenon also called paracellular BBBD [8]. However, a report on stroke [9], which is also associated with reversible brain edema, suggests a transcellular BBBD as the main mechanism of early edema formation in stroke. It remains unclear whether transcellular BBBD occurs in other diseases with reversible brain swelling, and which factors determine the reversibility of brain swelling.

Despite reversal of brain swelling structural changes, such as microhemorrhages, have been reported in survivors of CM [10, 11]. However, it remains unclear how and at what stage of the disease microhemorrhages occur. Reversibility of vasogenic brain edema and the occurrence of microhemorrhages can be assessed *in vivo* by MRI, but further invasive analyses in humans are currently limited. In addition, pathophysiological studies have been hampered by the lack of reliable animal models. Investigating the mechanisms underlying reversible edema is necessary to establish new therapeutic approaches and to eventually reduce permanent brain damage in CM. The development of a reliable animal model would thus represent a valuable tool to gain insight into these pathogenetic events.

Using serial *in vivo* MRI, we demonstrated for the first time the reproducible induction of reversible brain edema in a murine model of CM. We identified transcellular BBBD as an edema mechanism and showed that microhemorrhagic remnants occur in areas of higher BBBD. We further highlighted the translational potential of this experimental model by illustrating similar patterns of microhemorrhages in ECM and in human CM.

## Results

### Reversible brain swelling can be reliably induced in experimental cerebral malaria

To test whether the experimental cerebral malaria (ECM) model associates with reversible brain swelling, we performed serial MRI on C57Bl/6 mice infected with a low number (1000) of *Plasmodium berghei* (Pb ANKA) sporozoites. We examined BBBD and edema evolvement in all mice as soon as the first animal developed ECM and followed further progress of disease that led to either recovery or death. Non-surviving mice showed imaging characteristics on MRI that we previously described, including BBBD with rostral predominance (Fig. 1A) [12]. Notably, five out of twenty mice survived. These surviving mice also exhibited BBBD, albeit less pronounced (Fig. 1B). It is noteworthy that similar MRI findings have been demonstrated in 55% of *Pb* ANKA infected mice treated with a glutamine-antagonist [13]. However, the lack of predictive markers to identify edema reversibility at the acute stage of disease has hampered investigation of pathogenetic mechanisms associated with reversible versus irreversible edema. To circumvent this limitation, we set out to develop a model in which reversible edema could be reliably induced. We reasoned that partial protection of mice could be induced by immunization with attenuated sporozoites. A previous study demonstrated complete protection from ECM after single vaccination, with 100% survival, but no sterile protection against malaria infection [14]. We therefore investigated whether reversible edema occurred in mice after single vaccination with radiation-attenuated sporozoites (RAS). In our set up, all twenty wildtype mice after a single vaccination with RAS survived a subsequent challenge with sporozoites. Thirteen of these mice (65%) showed reversible BBBD (Fig. 1C). The remaining seven vaccinated mice did not show alterations in brain on MRI signal (five of these developed parasitemia). In comparison, only 25% of non-immunized wildtype survived survived after sporozoite challenge (Fig. 1D). Thus, a single vaccination with RAS is protective against fatal ECM but infected mice often develop reversible edema, a disease pattern which is only sporadically observed after infection of non-immunized wildtype mice.

**Fig. 1:**
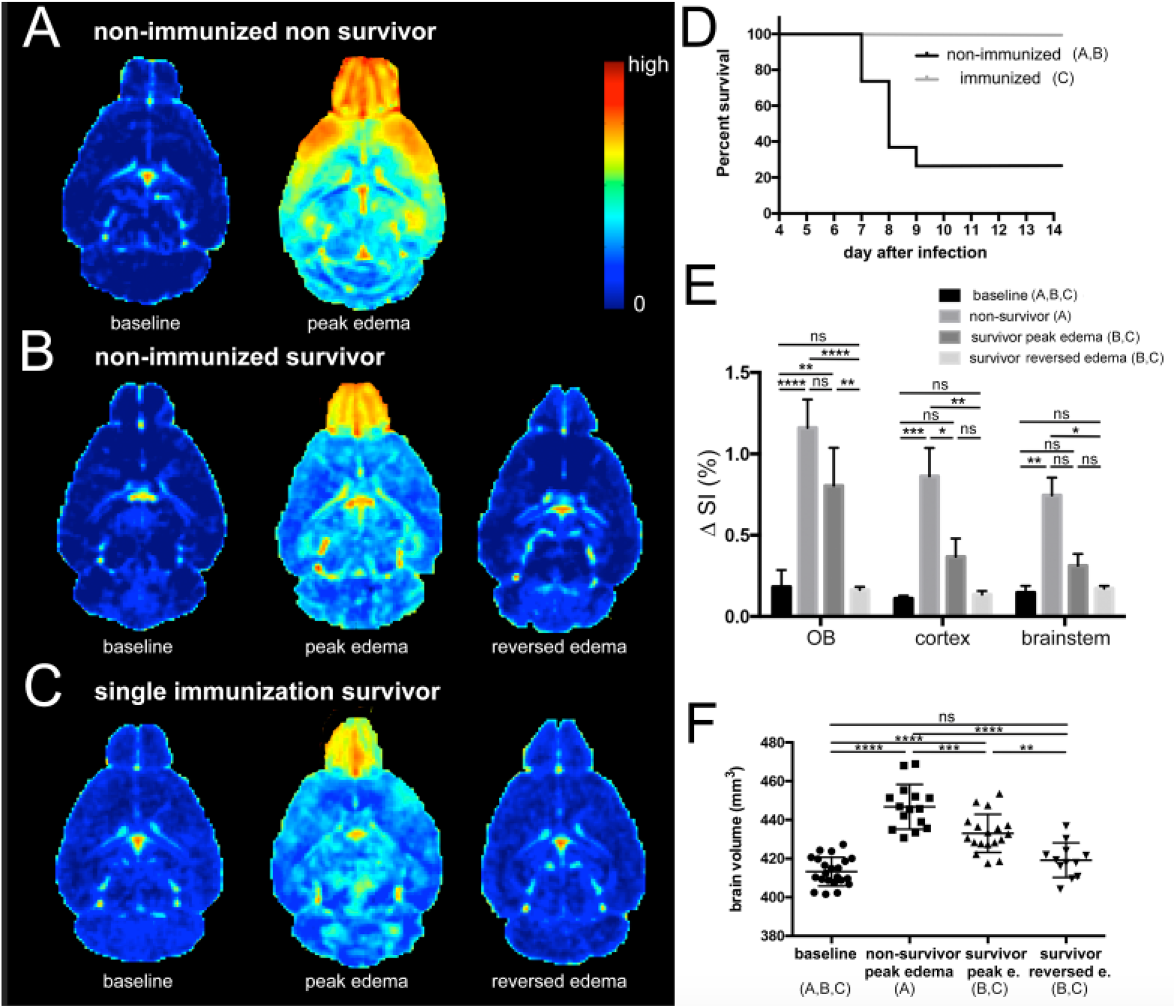
Reversible brain swelling in experimental cerebral malaria. A,B,C Subtraction images of T1-weighted images are displayed to illustrate the degree and spatial distribution of blood-brain barrier disruption (BBBD) in surviving and non-surviving mice. D, Survival curves of wildtype infected mice and mice after single vaccination. (A n=15, B n=5; C n=13) E, The difference in signal intensity (ΔSI) as a measure of BBBD is present in the olfactory bulb, cortex and brainstem in all groups. Most pronounced signal alterations are seen in the olfactory bulb. In cortex and brainstem less BBBD is seen in survivors. After edema resolves BBB, normalizes and returns to baseline values. F, The degree of brain swelling is shown at baseline, in non-survivors and survivors at the acute stage and after edema has reversed in surviving mice. Significance levels were tested with one-way ANOVA.

At peak edema survivors showed significantly less BBBD and significantly less brain volume increase compared to non-surviving mice (Fig. 1E, F). Overall, edema lasted one to three days before it reversed (Suppl. Fig. 1) and parasitemia increased at a slower rate in surviving mice compared to non-surviving mice (Suppl. Fig. 2). Peak edema in survivors was delayed by one day (9.3± 0.8 days post-infection) compared to non-survivors (8.1± 0.6 days post-infection). Survivors exhibited slower parasite growth rates and less fulminant edema, confirming that priming of the immune system and host-controlled parasite growth by single vaccination provides protection from severe disease [14-16]. At day 13, the last imaging time point, brain swelling had reversed and returned to baseline volumes (Fig. 1F).

### Transcellular blood brain barrier disruption mediates reversible brain edema

The single RAS-immunization model allowed us to study not only cerebral changes after edema reversal, but also for the first time its acute stage. We therefore addressed the mechanisms leading to the development of BBBD and its reversal by analyzing ultrastructural changes underlying these processes. We focused on the endothelium of small brain vessels using electron microscopy, which allows the discrimination between paracellular and transcellular BBBD.[9] While the former is triggered by ruptured tight junctions between endothelial cells, the latter is characterized by an increase of intra-endothelial vesicles transporting fluid from the luminal side to the brain parenchyma. In healthy control mice, intact TJs and few intracellular vesicles were apparent, both characteristics of an intact BBB (Fig. 2A). In non-survivors, 70% of TJs remained intact, while the number of intra-endothelial vesicles increased by 1,600% compared to healthy controls (Fig. 2A). Dissolution of the basal lamina was also seen (Suppl. Fig. 3A, B). In survivors at the acute stage of edema, TJs and basal lamina remained fully intact, while intracellular vesicles increased by 1,100% compared to healthy controls (Fig. 2A, Suppl. Fig. 3B). Upon the reversal of edema, the number of vesicles decreased by almost a half - to 700% of the healthy control level, while 97% of TJs remained intact (Fig. 2A), and disrupted TJs were seen in 3% of acquired images. No dissolution of the basal lamina was observed (Supplemental Fig. 3B). In order to test if vesicles serve as vehicle for inflammatory agents e.g. fluid and proteins to reach the parenchymal side, we injected DNP-albumin, and visualized it with immunogold particles (Fig. 2B). In healthy mice with no edema most DNP-albumin remained intraluminal, while it was visible within vesicles and on the parenchymal side in mice with edema (Fig. 2B).

**Fig. 2:**
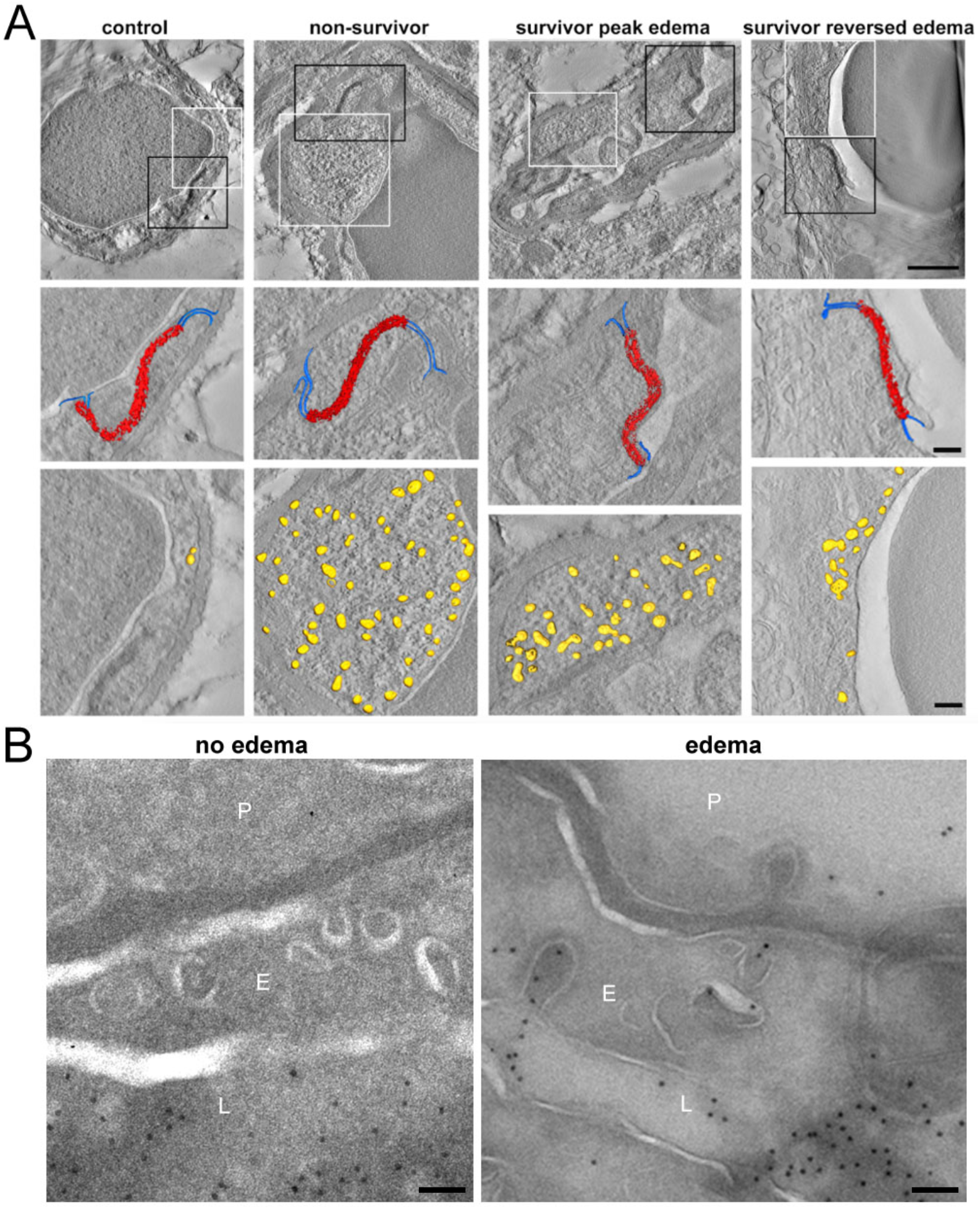
Reversible edema is induced by transcellular blood brain barrier disruption. A, Tomographic slices of example vessels are displayed with 3D models projected on the slice. Black squares are magnified below to demonstrate intact tight junctions (red) and adjacent adherence junctions (blue). White squares are magnified below and illustrate the increased number of vesicles in the endothelium (yellow), occurring during transcellular blood-brain barrier disruption. Scale bar first row 1 µm, scale bar second and third rows 200 nm. B, DNP-Gold labelled Tokuyasu sections of mice injected with DNP-albumin. In a control mouse without edema, DNP-albumin remains intraluminal. (first images). In an ECM mouse with edema, DNP-albumin reaches the parenchymal side via vesicles (second image). L=lumen, E=endothelium, P=parenchyma. Scale bar 100nm.

### Microvascular damage occurs after edema reversal

Microhemorrhages were visible by MRI in areas of most severe BBBD. These were predominantly located in the olfactory bulb and were more numerous in non-survivors (Fig. 3A). In survivors at peak of disease significantly less microhemorrhages were apparent, compared to non-survivors. Even though BBBD and edema reversed in survivors, microhemorrhages remained visible and surprisingly significantly increased after edema had reversed (Fig. 3A, B). This indicates that endothelial damage is severe in non-survivors, leading to immediate microhemorrhages. In survivors however, damage occurs after peak edema, when brain swelling reverses and blood flow normalizes. Impaired vessels may not withstand the increased pressure, thereby leading to microhemorrhages.

**Fig. 3:**
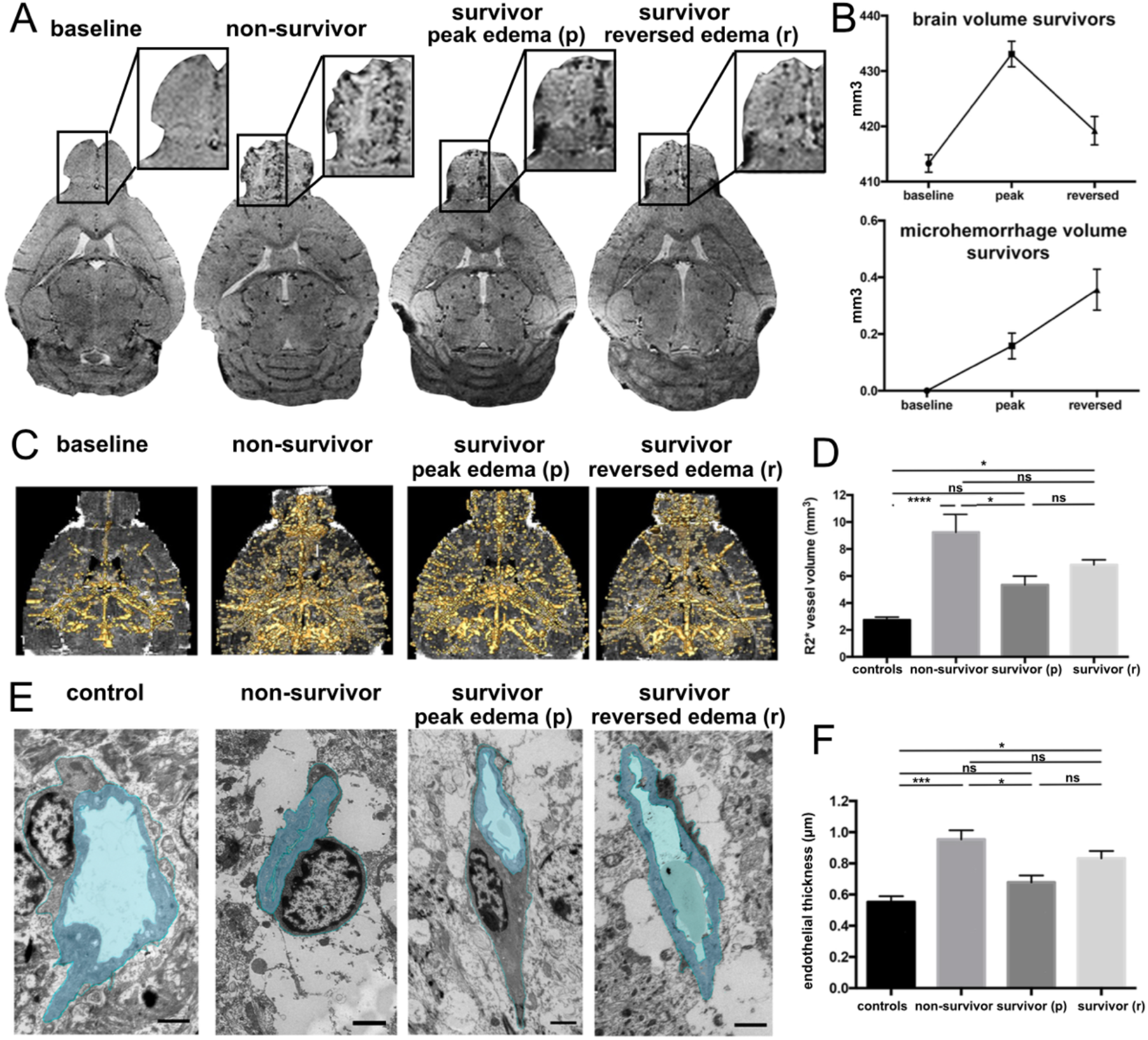
Reperfusion injury on the microscopic and ultrastructural level. A, Exemplary T2*w images are displayed. At baseline no microhemorrages are apparent (arrows). In severe disease a high microhemorrhage load is seen, mainly in the olfactory bulb (OB). In survivors microhemorrhages increase after edema has resolved. B, Graphs illustrate the increase of microhemorrhage volume while brain volume reverses. C, Segmented vessels (gold) on R2* datasets are displayed. The segmentation delineates vessel lumen and vessel wall. In non-survivors vessel volume is highest. In survivors it increases after edema reverses. D, R2* vessel volume throughout the groups is presented. (n=5-13). E, Ultrastructural endothelial changes on TEM images are illustrated by one example of a vessel from each group. In healthy mice the endothelium is thin with an open lumen. Astrocyte endfeet cover the endothelium. In non-survivors the endothelium is thicker, the lumen collapsed and surrounded by swollen astrocyte endfeet. Survivors in the acute stage of disease show a slight increase in endothial thickness, but also swollen astrocyte endfeet. Astrocyte endfeet swelling decreases after edema has reversed, but endothelial thickness slightly increases. F, Average endothelial thickness in all groups is displayed (n=3-5); Significance levels were tested with one-way ANOVA.

Interestingly, surviving mice that exhibited larger brain volumes in the acute stage of disease, developed significantly more microhemorrhages after edema had resolved, showing that initial disease severity correlates with the degree of microhemorrhages after recovery (Fig. 4A).

**Fig. 4:**
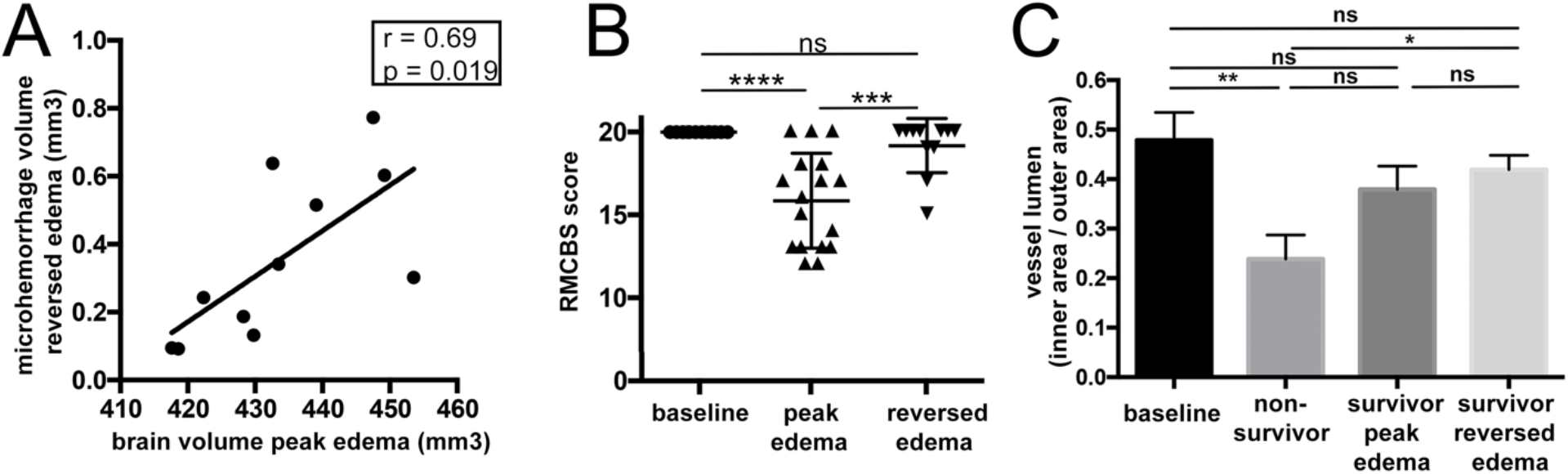
Correlation and time course of reperfusion injury and behavioral changes. A, Microhemorrhage volume correlates with the degree of brain swelling at the acute stage of the disease. B, Behavioral changes were assessed with the Rapid Murine Coma and Behavioral Scale (RMCBS) score. At baseline all mice displayed a healthy score of 20. Survivors showed altered behavior during the acute stage of disease but returned back to normal in most mice. C, The ratio of inner area of the endothelium versus the outer area of the endothelium is displayed as a measure of collapse of lumen. Significance levels were tested with one-way ANOVA. Spearman’s analyzes were used for correlation analyses.

In order to investigate the association between imaging features and behavioral alterations, the rapid murine coma and behavioral scale (RMCBS) score was employed. 30% of surviving mice did not reach a healthy baseline score of 20 after edema had resolved. The remaining 70% reached a baseline score of 20 despite exhibiting microhemorrhages (Fig. 4B). To further analyze if the vasculature itself shows impairment, we examined the vasculature in affected brain volumes at microscopic and ultrastructural levels. On MRI-derived R2* maps, which quantify susceptibility signal of deoxygenized blood on MR images, we assessed the vascular volume *in vivo* over time at a resolution of 50µm. Interestingly, vessel volume within R2* maps in survivors increased when edema had reversed, corroborating that injurious processes continue developing further (Fig 3C, D). Also at the ultrastructural level endothelial alterations matched the observed *in vivo* changes detected by R2* maps. Even though vessel lumen normalized after edema had reversed (Fig. 4C), the endothelial perimeter increased in survivors, indicating endothelial remodeling and consecutive increased vascular reactivity that promotes peripheral resistance (Fig. 3 E, F). Altogether, these findings provide evidence that microvascular impairment occurs after edema reversal.

### Translational potential of the experimental model

To investigate the relevance of our model to human CM, we analyzed MR datasets that included susceptibility-weighted imaging (SWI), a sensitive MRI sequence to detect microhemorrhages, from both pediatric and adult Indian CM patients admitted at Ispat General Hospital in Rourkela, India as part of a study described elsewhere [17, 18]. In this subgroup, similar distributions of brain volume were apparent with higher normalized brain volumes in pediatric CM compared to adult CM (Figure 5A) [18]. CM patients with microhemorrhages showed a wide range of brain volumes (Figure 5A). As only one SWI dataset was acquired in most patients, we could not assess the temporal course of microhemorrhage occurrence. The presence of microhemorrhages did not differ in adult and pediatric CM, with 50% in fatal CM (1 out of 2 pediatric CM patients as well as 1 out of 2 CM adult patients), 33% in non-fatal pediatric CM (3 out of 9 patients) and 36% in non-fatal adult CM (5 out of 14 patients) (Supplemental Table 1). Similar to the experimental model, microhemorrhages are more frequent in fatal compared to non-fatal disease during the acute stage of the disease. The anatomic predilection site of microhemorrhages in the brain was not the olfactory bulb but the grey and white matter junction (Figure 5B), as well as the corpus callosum, basal ganglia and cerebellum. Despite different microhemorrhage location in the mouse model and human CM, microhemorrhages occur during both pediatric and adult CM and may be due to reperfusion injury.

**Figure 5:**
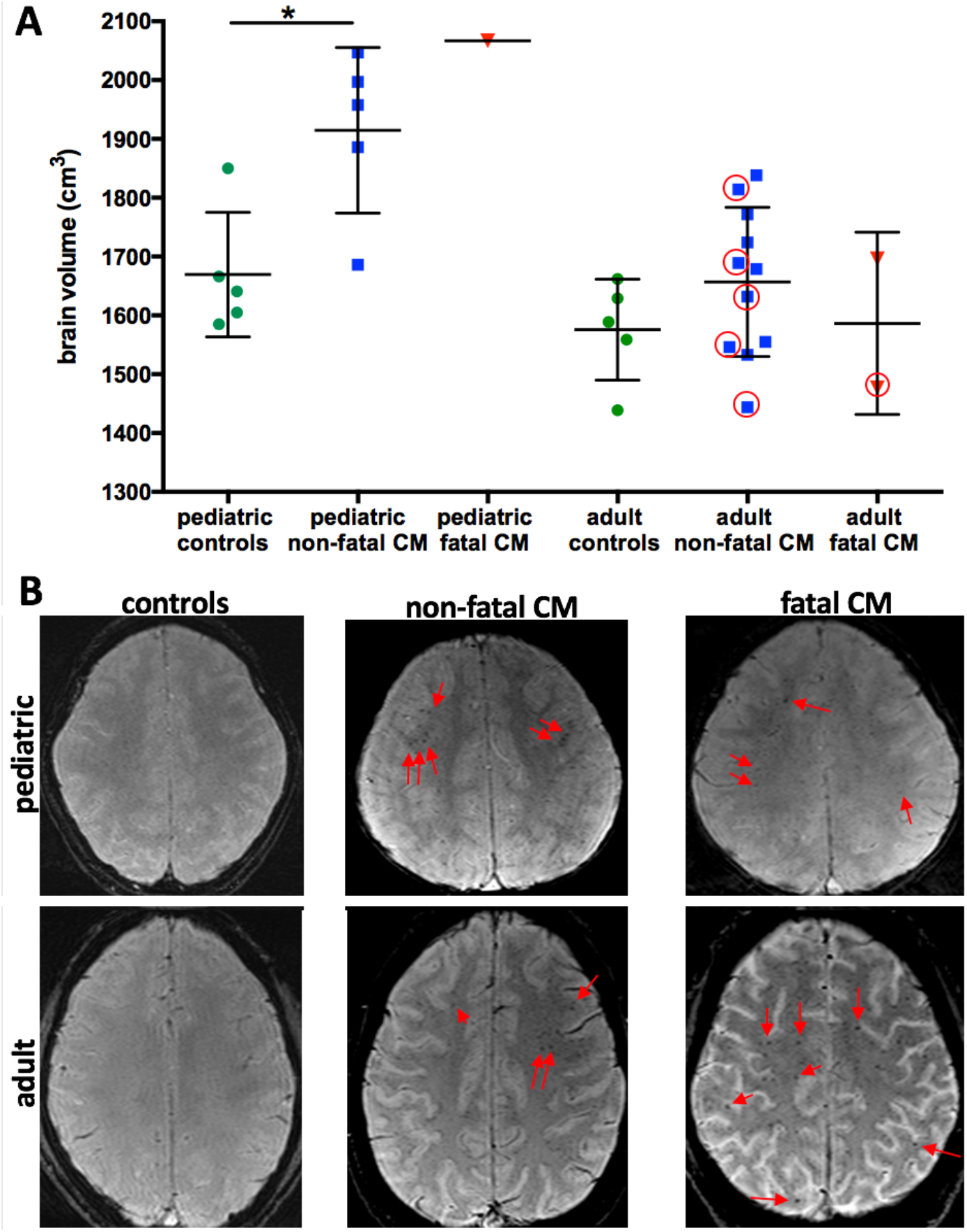
Brain swelling and occurrence of microhemorrhages in pediatric and adult CM. A. Normalized brain volume in children was significantly increased compared to age-matched controls and more pronounced in children compared to adults. In adults no significant increase compared to healthy volunteers was noted, despite a spread in brain volume in non-fatal CM and fatal CM. Red circles label data points of patients with successful brain volume calculation and presence of microhemorrhages. In this cross-sectional dataset brain volumes of patients with microhemorrhages show a wide range. Significance levels were tested with one-way ANOVA. B. Exemplary susceptibility-weighted images are shown. In controls no microhemorrhages are visible. In non-fatal and fatal diseas red arrows point to exemplary microhemorrhages at the subcortical white matter.

## Discussion

In this study, we demonstrated that i) transient, reversible brain swelling can reliably be induced in ECM, ii) that transcellular BBBD is an important mechanism of edema development in ECM and iii) that permanent damage remains in areas of most severe BBBD. We further highlighted the translational potential of the model by showing similarities of the experimental model with clinical findings.

Developing this novel model allowed us to investigate for the first time the pathogenic events during the acute phase of the disease leading to reversible vasogenic edema. We identified transcellular BBBD without disrupted tight junctions (TJ) as a mechanism of brain swelling in ECM. Upregulation of intra-endothelial vesicles is the initial response of brain endothelium to ischemia/reperfusion and to inflammation but has been largely overlooked as a step of early BBBD [9, 19-21]. Although the number of intra-endothelial vesicles increased, intact TJs indicate that endothelial cells keep their lining, without pulling TJ complexes apart or being destroyed by proteinases, thus enabling edema reversal [22]. This is in line with our observations in fatal ECM, as well as previous findings in both ECM and CM, showing disrupted TJs and higher numbers of intra-endothelial vesicles associated with irreversible edema [5, 23-25]. By analyzing both survivors and non-survivors for the first time and by performing tracer experiments, we now extend previous reports and identify endothelial vesicles and thus transcellular BBBD as a contributing mechanism to brain swelling in our model of ECM. In other diseases such as stroke, transcellular BBBD precedes TJ disruption, which occurs in later phases of BBBD and represents potentially more severe damage [9, 26]. It is therefore conceivable that transcellular BBBD is not only responsible for reversible vasogenic edema in CM but may also be involved in reversible brain edema in general.

Despite normalization of paracellular BBBD and edema reversal, a delayed occurrence of microhemorrhages and vascular remodeling was observed in animals who survived ECM. Small hemorrhages have been linked to severe transcellular BBBD after reperfusion in an experimental stroke model [27]. In our study, microhemorrhages mainly occurred in a delayed fashion in the olfactory bulb, the region of most pronounced BBBD. Remarkably, in a model of hypoxia/reperfusion, which mimics high altitude cerebral edema, mice exhibited a similar temporal course of microhemorrhages with a strong increase after reoxygenation in the same anatomic areas. [28] The location of microhemorrhages in human CM differed from the experimental model and are seen predominantly in the corpus callosum, the subcortical white matter and the basal ganglia. However, they developed at the same locations as in other human disease with hypoxia/reperfusion, such as high-altitude edema [29, 30]. These similarities suggest that *(i)* hypoxia is an important factor in ECM/CM pathogenesis; *(ii)* reperfusion and reoxygenation may lead to microhemorrhages; and *(iii)* they indicate prior vascular damage during acute disease. The importance of hypoxia during ECM pathogenesis and during CM has been described previously [18, 31, 32], and the observed endothelial swelling may also be induced by hypoxia [33].

We expand these findings and further demonstrate that reperfusion injury occurs in a similar fashion to other diseases that involve hypoxia/reperfusion. Interestingly, a review article about CM from 1997 postulated that microhemorrhages may be induced by reperfusion injury due to their distinct pathological features [34]. Indeed, the unregulated restoration of flow in a damaged cerebral microvessel previously filled with sequestered parasitized red blood cells and host monocytes may cause its rupture, forcing the contents out into the brain parenchyma, and a subsequent leakage of non-parasitized erythrocytes [34].

Since we show that microhemorrhages in ECM/CM survivors occur when brain swelling reverses, microhemorrhages are highly likely to be caused by reperfusion injury and mark areas that are most severely damaged during acute disease. These cerebral microhemorrhagic ‘scars’ can also be detected by MRI on long-term follow-up scans in diseases with reversible brain swelling [35]. The number and intensity of microhemorrhages may thus be used as an indicator of initial disease severity in CM. It is noteworthy that microhemorrhages may not be clinically silent. They have been associated with long-term neurological impairment in patients in several diseases, including diffuse axonal injury, small vessel disease and cognitive decline in dementia, and may represent markers for the degree of neurocognitive impairment [36-38]. Neurological sequelae are known to occur after CM [3] and could potentially be linked to microhemorrhages. They may thus represent a powerful marker to predict potential neurocognitive impairment and warrant further clinical investigation.

Taken together, our results suggest a potential association between the degree of initial transcellular BBBD, consecutive brain swelling and reperfusion injury. This association indicates that adjuvant drug treatment targeting transcytosis could reduce reperfusion injury and protect from long term neurocognitive impairment.

## Material and Methods

### Ethics statement

Mouse model: All animal experiments were performed according to FELASA category B and ARRIVE guidelines and approved by the local German authorities in Karlsruhe. Patients: Ethical approval was obtained from The Indian Council of Medical Research as well as from the institutional review boards of Ispat General Hospital, New York University School of Medicine and the London School of Hygiene and Tropical Medicine. Because CM patients were comatose, written informed consent was obtained from the families of all patients before enrollment in the study as previously described [17, 18].

### Murine malaria model

ECM was induced with the *Plasmodium berghei* ANKA (*Pb* ANKA) parasite in inbred female C57BL/6J mice (Janvier Labs, France). *Pb* ANKA sporozoites (SPZ) were isolated by dissection of salivary glands from female *Anopheles stephensi* mosquitoes at day 18-21 post infection. Infections of 6-8-week-old female mice were performed by intravenous (i.v.) injections of 1000 SPZ in a total volume of 100µl sterile PBS (n=12). In a second group the same infections in 12-14-week-old mice were carried out in naïve mice (n=8) and single vaccinated mice (n=20) (Heiss et al., 2018). For immunization with radiation-attenuated sporozoites (RAS), SPZ were treated by exposure to 150 Gy of γ-radiation (^137^Cesium source, University Hospital Heidelberg, Germany) and were then injected into mice at a dose of 3×10^4^ RAS. Brains of three healthy female C57BL/6J mice were used as controls for tissue analysis. Parasitemia was assessed starting at day 4 after infection. For clinical evaluation, malaria-infected mice were assessed for ten parameters of cerebral symptoms according to the Rapid-Murine-Coma-and-Behavioral-Scale (RMCBS)[39]. The RMCBS testing was performed daily starting at day 6 after infection.

### Experimental MRI protocol

MRI was performed on a 9.4 T small animal scanner (BioSpec 94/20 USR, Bruker Biospin GbmH, Ettlingen, Germany) using a volume resonator for transmission and a 4-channel-phased-array surface receiver coil. Anesthesia was induced per inhalation using 2 % and maintained with 1-1.5 % isoflurane. Mice were placed prone in fixed position monitoring body temperature and respiration. Starting at day 6 after infection mice were screened for edema by using a 2D T2-weighted sequence (repetition time/ echo time (TR/TE)=2000/22ms, slices=12, slice thickness=0.7mm). The baseline MR imaging protocol before infection and the MR imaging protocol at occurrence of edema and at day 14 after infection included 3D T1-weighted imaging (TR/TE=5/1.9ms, flip angle (FA)=8.5°, 156µm isotropic resolution) before and after injection of 0.3 mmol/kg Gd-DTPA and T2*-weighted flow compensated gradient echo imaging (TR/TE=50/18ms, FA=12°, 80µm isotropic resolution) or T2* multi gradient echo imaging (TR=50ms, TE=3.5ms-32.2ms with increments of 5.7ms, FA=14°, 100µm isotropic resolution).

### Electron microscopy of mouse tissue

For morphological analysis mice were transcardially perfused with 0.9% saline after the last MRI scan. Brains were removed and fixed in 2% glutaraldyhyde in saline overnight at +4 degrees. OB tissue was cut in cubes and postfixed with 2% glutaraldehyde + 2% paraformaldehyde in 100mM cacodylate buffer, followed by fixation in 1% Osmium tetroxide in cacodylate buffer and contrasting with 1% uranylacetate in water. Samples were dehydrated through immersion in a series of increasing percentage of acetone and embedded in Spurr’s resin (Serva, Heidelberg, Germany). Sectioning was done on a Leica UC6 microtome (Leica Microsystems, Vienna, Austria) and 70nm sections were collected on Formvar-coated, copper mesh grids and imaged on a JEOL JEM-1400 electron microscope (JEOL, Tokyo) operating at 80 kV and equipped with a 4K TemCam F416 (Tietz Video and Image Processing Systems GmBH, Gautig). For tomography 350nm thick sections were placed on formvar coated slot grids. Tilt series over a ±60° range were recorded in a Tecnai F20 EM (FEI, Eindhoven, The Netherlands) operating at 200kV using the SerialEM software package (Mastronarde 2005) on FEI Eagle 4K x 4K CCD camera at a magnification of 9.5kx resulting in 2.24 nm pixel size. Tilted images were aligned by cross-correlation procedure. The tomograms were generated by weighted-back-projection algorithm using IMOD processing packages [40].

For tracer experiments with Dinitrophenyl (DNP)-albumin conjugate (Sigma-Aldrich), 50mg DNP-Albumin were injected i.v. after the last MRI scan and circulated for 30 minutes. Brains were removed and fixed in 4% paraformaldehyde, 0.016% glutaraldyhyde in saline overnight at +4 degrees and then cut into 150µm sections. After a 2-hour fixation step with 4% paraformaldehyde and PHEM buffer, small tissue cubes were infiltrated with gelatine of increasing concentrations (1%, 6%, and 12%). After incubation overnight in 12% gelatine, tissue cubes were incubated with 2.3M sucrose overnight and then frozen in liquid nitrogen. Sectioning was done on a Leica UC6 cryo-microtome (Leica Microsystems, Vienna, Austria). Sections were immunolabelled with a rabbit anti-DNP antibody (ABnostics, Dossenheim, Germany) at a dilution of 1:50 and in a second step with goat anti-rabbit antibody coupled to 10 nm protein A gold (CMC university medical center Utrecht, Netherlands) diluted 1:50.

### Image analysis of experimental data

Image processing was undertaken in Amira 5.4 (FEI, Visualization Sciences Group). Edema was graded on 3D T2-weighted images into mild (1), moderate (2) and severe (3) as previously described [12]. Blood-brain barrier permeability (BBBD) was assessed by contrast-enhanced 3D gradient echo T1-w imaging. 3D non-enhanced T1w images were subtracted from contrast enhanced T1-weighted images. In pre- and post-contrast 3D T1-weighted images, Gibbs ringing was suppressed, and signal-to-noise-ratio enhanced using a 3D spatial Gaussian low-pass filter with a resulting effective isotropic resolution of 280µm. In case of significant motion between the sequences, images were motion corrected using a custom-made MATLAB code for rigid body registration. First the difference images were evaluated for pathological enhancement by visual inspection. Second, the relative signal enhancement ΔSI (%) in different regions-of-interest (ROI) was quantified as: ΔSI (%) = [(SI_post contrast_ – SI_pre contrast_) / SI_pre contrast_ ] x 100%. ROIs were placed after anatomical delineation manually into the following structures: 1) olfactory bulb (OB)+rostral-migratory-stream (RMS), 2) dorsal-migratory-stream (DMS), 3) external capsule (EC), 4) cortex, 5) basal ganglia, 6) thalamus and 7) brainstem (BS) according to the Allen Brain Atlas [41]. RMS and DMS are only visible during disease, as in healthy mice they display the same signal intensity as the surrounding tissue. Therefore, ROIs were drawn at the estimated location of the structures on the scans without signal alterations in these areas. Microhemorrhage volume was semiautomatically segmented by manual region growing using a threshold-based presegmentation on T2w* datasets.

Vessel volume was semiautomatically segmented on R2* maps, which were assessed by voxel-wise monoexponential fitting of the T2* signal decay using the freely available relaxometry tool (https://github.com/neuroimaging-mug/relaxometry). Tight junctions and endothelial vesicles were semiautomatically segmented with Amira. Endothelial measurements were performed on EM images of vessel cross sections using ImageJ (version 1.49s). [42] Endothelial thickness was defined as a ratio of the area of vessel lumen in cross section to that of the whole vessel. Endothelial perimeter was measured along eight equally spaced lines, expanding radially from the center of the vessel lumen.

### Study site and patients

The study was carried out at Ispat General Hospital (IGH) in Rourkela, in the state of Odisha, India, from October 2013 to November 2019. [17, 18] Patients with cerebral malaria (CM) that underwent susceptibility-weighted imaging were included into the analysis (total n=27, fatal n=4, non-fatal n=23). All CM patients satisfied a strict definition of CM according to the modified World Health Organization criteria. CM patients with coma (defined as a Glasgow coma score [GCS] of 9 out of 15 for adults and a Blantyre coma score [BCS] of 2 for young children) after correction of hypoglycemia (2.2 mmol/liter) and infected with *Plasmodium falciparum* (detected by rapid diagnostic test and confirmed by the presence of asexual forms of the parasite in a peripheral blood smear) fulfilled inclusion criteria. Healthy subjects served as controls. The adult controls were imaged at the same MRI at IGH. Age-matched pediatric controls were retrospectively recruited at Heidelberg University Hospital.

### Human MRI protocol and image analysis

Imaging was performed using a 1.5-Tesla (T) Siemens Symphony MRI scanner (Siemens AG, Erlangen, Germany). Scanning was carried out within 10 h of admission. The MRI sequences included axial T1-weighted (TE/TR=7.7ms/500ms, slice thickness=5mm), T2-weighted (TE/TR=99ms/4000ms, slice thickness=5mm), and susceptibility-weighted imaging (TE/TR=40ms/50ms, slice thickness=2mm, flip angle=12 degrees). Normalized brain volume of T1-weighted images was calculated using SIENAX, which is part of the FSL toolbox (Smith et al., 2004). The occurrence of brain swelling and microhemorrhages was visually assessed on T2-weighted and susceptibility-weighted images, respectively.

### Statistics

Data are shown as mean ± standard error of mean (SEM) Statistical analyses was performed in PRISM (GraphPad, version 7, La Jolla, CA). To compare two groups, unpaired, two tailed student’s t-tests were used (e.g. parasitemia non-survivor, survivor). To compare more than two experimental groups (baseline/control, non-survivor, survivor) one-way analysis of variance (ANOVA) with multiple comparisons was performed. Spearman’s analyzes were used for correlation analyses. p values ≤ 0.05 were considered statistically significant.

## Acknowledgements

The expert technical assistance of Miriam Reinig, Stephanie Gold, Akshaya Mohanty and Nakul C. Khatua is gratefully acknowledged. We thank Charlotta Funaya, Stefan Hillmer and the Electron Microscopy Core Facility of Heidelberg University for their valuable support, as well as the Director in Charge and the clinical staff of Ispat General Hospital in Rourkela for their help and dedication. We further thank Gareth G Griffiths for his very helpful comments and proofreading of the manuscript.

Angelika Hoffmann was supported by the Olympia Morata Fellowship of the Medical Faculty of the University of Heidelberg, a sCDF fellowship of the MMPU partnership unit and a clinical leave stipend from the German Centre for Infection Research (Deutsches Zentrum fuer Infektionsforschung, DZIF. Jessica Jin and Chi Ho Wai are supported by an MD fellowship from the German Centre for Infection Research (Deutsches Zentrum fuer Infektionsforschung, DZIF). Sanjib Mohanty, Samuel C. Wassmer, and Praveen K. Sahu are supported by the National Institute of Allergy and Infectious Diseases of the National Institutes of Health (NIH) under award number U19AI089676. Samuel C. Wassmer is also supported by a Research Project Grant from the UK Medical Research Council (award number MR/S009450/1) and an NIH R21 grant together with Angelika Hoffmann (award number R21AI142472). The content is solely the responsibility of the authors and does not necessarily represent the official views of the NIH. Michael Lanzer and Friedrich Frischknecht are supported by the Deutsche Forschungsgemeinschaft under the SFB1129.

## Author contributions

Study concepts/study design or data acquisition or data analysis/interpretation, all authors; manuscript drafting or manuscript revision for important intellectual content, all authors; approval of final version of submitted manuscript, all authors; agrees to ensure any questions related to the work are appropriately resolved, all authors; literature research, A.H., A.K.M., J.J., S.C.W., M.C.; experimental studies, A.H., J.J., J.P., A.K.M., C.H.W., M.F., M.C., M.A.B.; clinical studies, S.M., P.K.S., R.P., M.M., S.C.W.; statistical analysis, A.H., M.C. and manuscript editing, A.H., S.C.W., F.F., M.C., M.B.

## Supporting Material

**Suppl. Fig. 1:**
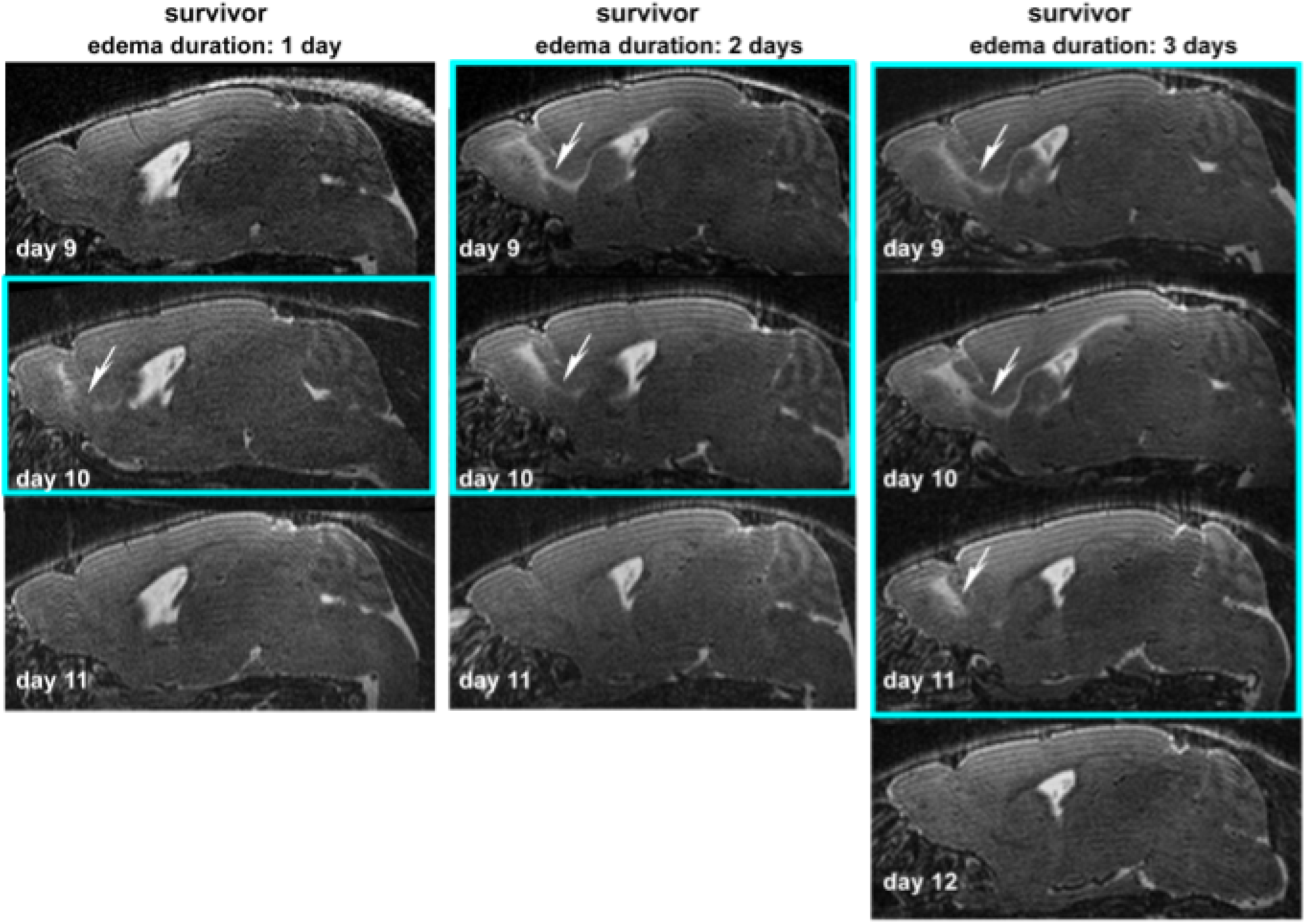
Edema duration. Exemplary T2w sagittal images of three surviving mice are displayed. Turquoise squares surround images with edema. Edema (white arrows), visible as increased (bright) T2w signal was evident for three days, if it had reached the dorsal migratory stream, or lasted for one to two days if mainly the olfactory bulb and rostral migratory stream were affected.

**Suppl. Fig. 2:**
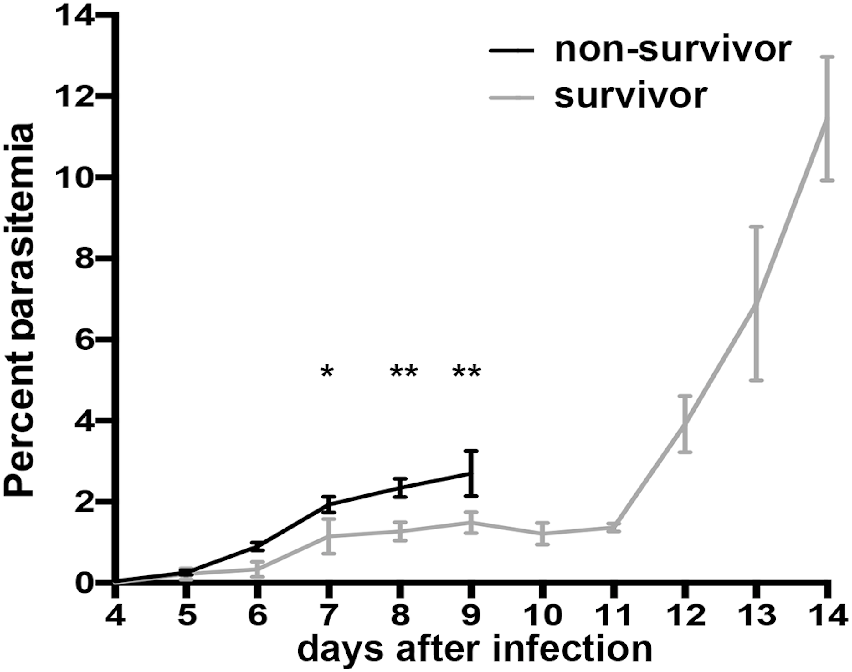
Parasitemia course. Parasitemia increases slower in surviving mice and is significantly different to non-survivors from day 7 to day 9. * indicates significance levels as calculated by the unpaired, two tailed student’s t-test.

**Suppl. Fig. 3:**
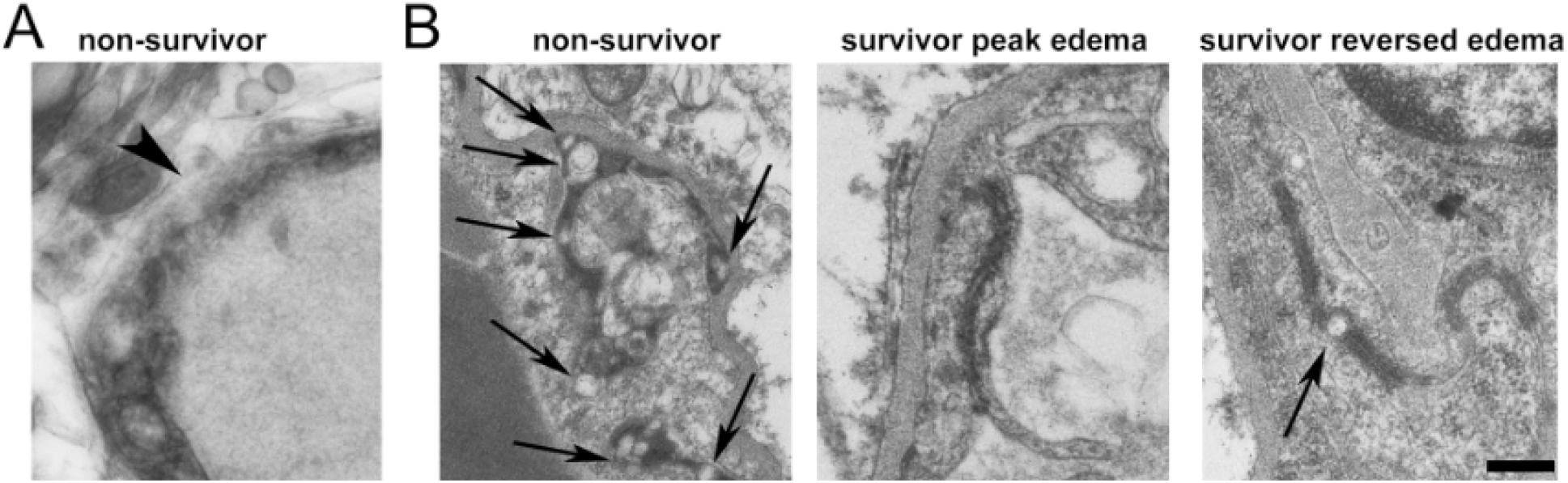
Basal lamina and tight junctions. A, Dissolvement of the basal lamina in an example vessel from a non-surviving mouse is shown (arrowhead). B, Abnormal tight junction with gaps (arrows) are seen in non-survivors and rarely in mice after edema had reversed, but not in surviving mice at peak edema. Scale bar 200 nm.

**Supplemental Table 1:** Number of CM patients with microhemorrhages

|  | Pediatric CM non-fatal | Adult CM non-fatal | Pediatric CM fatal | Adult CM fatal |
| --- | --- | --- | --- | --- |
| Microhemorrhages present | 3/9 (33%) | 5/14 (36%) | 1/2 (50%) | 1/2 (50%) |
| Microhemorrhages absent | 6/9 (66%) | 9/14 (64%) | 1/2 (50%) | 1/2 (50%) |

## References

1. WHO. World Malaria Report 2016. 2016.

2. Marsh K, Forster D, Waruiru C, Mwangi I, Winstanley M, Marsh V, et al. Indicators of life-threatening malaria in African children. The New England journal of medicine. 1995;332(21):1399–404. doi: 10.1056/NEJM199505253322102. PubMed PMID: 7723795.

3. Boivin MJ, Bangirana P, Byarugaba J, Opoka RO, Idro R, Jurek AM, et al. Cognitive impairment after cerebral malaria in children: a prospective study. Pediatrics. 2007;119(2):e360–6. doi: 10.1542/peds.2006-2027. PubMed PMID: 17224457; PubMed Central PMCID: PMCPMC2743741.

4. Seydel KB, Milner DA Jr.,, Kamiza SB, Molyneux ME, Taylor TE. The distribution and intensity of parasite sequestration in comatose Malawian children. The Journal of infectious diseases. 2006;194(2):208-5. Epub 2006/06/17. doi: 10.1086/505078. PubMed PMID: 16779727; PubMed Central PMCID: PMC1515074.

5. Strangward P, Haley MJ, Shaw TN, Schwartz JM, Greig R, Mironov A, et al. A quantitative brain map of experimental cerebral malaria pathology. PLoS pathogens. 2017;13(3):e1006267. doi: 10.1371/journal.ppat.1006267. PubMed PMID: 28273147; PubMed Central PMCID: PMCPMC5358898.

6. Nishanth G, Schluter D. Blood-Brain Barrier in Cerebral Malaria: Pathogenesis and Therapeutic Intervention. Trends in parasitology. 2019. doi: 10.1016/j.pt.2019.04.010. PubMed PMID: 31147271.

7. Stokum JA, Gerzanich V, Simard JM. Molecular pathophysiology of cerebral edema. Journal of cerebral blood flow and metabolism : official journal of the International Society of Cerebral Blood Flow and Metabolism. 2016;36(3):513–38. doi: 10.1177/0271678X15617172. PubMed PMID: 26661240; PubMed Central PMCID: PMCPMC4776312.

8. Yang Y, Rosenberg GA. Blood-brain barrier breakdown in acute and chronic cerebrovascular disease. Stroke. 2011;42(11):3323–8. doi: 10.1161/STROKEAHA.110.608257. PubMed PMID: 21940972; PubMed Central PMCID: PMCPMC3584169.

9. Knowland D, Arac A, Sekiguchi KJ, Hsu M, Lutz SE, Perrino J, et al. Stepwise recruitment of transcellular and paracellular pathways underlies blood-brain barrier breakdown in stroke. Neuron. 2014;82(3):603-17. Epub 2014/04/22. doi: 10.1016/j.neuron.2014.03.003. PubMed PMID: 24746419; PubMed Central PMCID: PMC4016169.

10. Nickerson JP, Tong KA, Raghavan R. Imaging cerebral malaria with a susceptibility-weighted MR sequence. AJNR American journal of neuroradiology. 2009;30(6):e85–6. doi: 10.3174/ajnr.A1568. PubMed PMID: 19321625.

11. Potchen MJ, Kampondeni SD, Seydel KB, Haacke EM, Sinyangwe SS, Mwenechanya M, et al. 1.5 Tesla Magnetic Resonance Imaging to Investigate Potential Etiologies of Brain Swelling in Pediatric Cerebral Malaria. Am J Trop Med Hyg. 2018;98(2):497–504. doi: 10.4269/ajtmh.17-0309. PubMed PMID: 29313473; PubMed Central PMCID: PMCPMC5929182.

12. Hoffmann A, Pfeil J, Alfonso J, Kurz FT, Sahm F, Heiland S, et al. Experimental Cerebral Malaria Spreads along the Rostral Migratory Stream. PLoS pathogens. 2016;12(3):e1005470. doi: 10.1371/journal.ppat.1005470. PubMed PMID: 26964100; PubMed Central PMCID: PMCPMC4786214.

13. Riggle BA, Sinharay S, Schreiber-Stainthorp W, Munasinghe JP, Maric D, Prchalova E, et al. MRI demonstrates glutamine antagonist-mediated reversal of cerebral malaria pathology in mice. Proceedings of the National Academy of Sciences of the United States of America. 2018;115(51):E12024–E33. doi: 10.1073/pnas.1812909115. PubMed PMID: 30514812; PubMed Central PMCID: PMCPMC6304986.

14. Heiss K, Maier MI, Hoffmann A, Frank R, Bendszus M, Mueller AK, et al. Protection from experimental cerebral malaria with a single intravenous or subcutaneous whole-parasite immunization. Scientific reports. 2018;8(1):3085. doi: 10.1038/s41598-018-21551-2. PubMed PMID: 29449638.

15. Ademolue TW, Awandare GA. Evaluating antidisease immunity to malaria and implications for vaccine design. Immunology. 2018;153(4):423–34. doi: 10.1111/imm.12877. PubMed PMID: 29211303; PubMed Central PMCID: PMCPMC5838420.

16. Khoury DS, Cromer D, Akter J, Sebina I, Elliott T, Thomas BS, et al. Host-mediated impairment of parasite maturation during blood-stage Plasmodium infection. Proceedings of the National Academy of Sciences of the United States of America. 2017;114(29):7701–6. doi: 10.1073/pnas.1618939114. PubMed PMID: 28673996; PubMed Central PMCID: PMCPMC5530648.

17. Mohanty S, Benjamin LA, Majhi M, Panda P, Kampondeni S, Sahu PK, et al. Magnetic Resonance Imaging of Cerebral Malaria Patients Reveals Distinct Pathogenetic Processes in Different Parts of the Brain. mSphere. 2017;2(3). doi: 10.1128/mSphere.00193-17. PubMed PMID: 28596990; PubMed Central PMCID: PMCPMC5463026.

18. Sahu PK, Hoffmann A, Majhi M, Pattnaik R, Patterson C, Mahanta KC, et al. Brain Magnetic Resonance Imaging Reveals Different Courses of Disease in Pediatric and Adult Cerebral Malaria. Clinical infectious diseases : an official publication of the Infectious Diseases Society of America. 2020. Epub 2020/12/16. doi: 10.1093/cid/ciaa1647. PubMed PMID: 33321516.

19. Ito U, Ohno K, Yamaguchi T, Takei H, Tomita H, Inaba Y. Effect of hypertension on blood-brain barrier. Change after restoration of blood flow in post-ischemic gerbil brains. An electronmicroscopic study. Stroke. 1980;11(6):606–11. PubMed PMID: 7210066.

20. Lossinsky AS, Shivers RR. Structural pathways for macromolecular and cellular transport across the blood-brain barrier during inflammatory conditions. Review. Histol Histopathol. 2004;19(2):535–64. doi: 10.14670/HH-19.535. PubMed PMID: 15024715.

21. Kang EJ, Major S, Jorks D, Reiffurth C, Offenhauser N, Friedman A, et al. Blood-brain barrier opening to large molecules does not imply blood-brain barrier opening to small ions. Neurobiol Dis. 2013;52:204–18. doi: 10.1016/j.nbd.2012.12.007. PubMed PMID: 23291193.

22. Lossinsky AS, Vorbrodt AW, Wisniewski HM. Scanning and transmission electron microscopic studies of microvascular pathology in the osmotically impaired blood-brain barrier. J Neurocytol. 1995;24(10):795–806. PubMed PMID: 8586999.

23. Ampawong S, Chaisri U, Viriyavejakul P, Nontprasert A, Grau GE, Pongponratn E. Electron microscopic features of brain edema in rodent cerebral malaria in relation to glial fibrillary acidic protein expression. Int J Clin Exp Pathol. 2014;7(5):2056–67. PubMed PMID: 24966914; PubMed Central PMCID: PMCPMC4069908.

24. Lackner P, Beer R, Helbok R, Broessner G, Engelhardt K, Brenneis C, et al. Scanning electron microscopy of the neuropathology of murine cerebral malaria. Malaria journal. 2006;5:116. doi: 10.1186/1475-2875-5-116. PubMed PMID: 17125519; PubMed Central PMCID: PMCPMC1676017.

25. MacPherson GG, Warrell MJ, White NJ, Looareesuwan S, Warrell DA. Human cerebral malaria. A quantitative ultrastructural analysis of parasitized erythrocyte sequestration. The American journal of pathology. 1985;119(3):385–401. PubMed PMID: 3893148; PubMed Central PMCID: PMCPMC1888001.

26. Nag S, Venugopalan R, Stewart DJ. Increased caveolin-1 expression precedes decreased expression of occludin and claudin-5 during blood-brain barrier breakdown. Acta neuropathologica. 2007;114(5):459–69. doi: 10.1007/s00401-007-0274-x. PubMed PMID: 17687559.

27. Haley MJ, Lawrence CB. The blood-brain barrier after stroke: Structural studies and the role of transcytotic vesicles. Journal of cerebral blood flow and metabolism : official journal of the International Society of Cerebral Blood Flow and Metabolism. 2017;37(2):456–70. doi: 10.1177/0271678X16629976. PubMed PMID: 26823471; PubMed Central PMCID: PMCPMC5322831.

28. Hoffmann A, Kunze R, Helluy X, Milford D, Heiland S, Bendszus M, et al. High-Field MRI Reveals a Drastic Increase of Hypoxia-Induced Microhemorrhages upon Tissue Reoxygenation in the Mouse Brain with Strong Predominance in the Olfactory Bulb. PloS one. 2016;11(2):e0148441. doi: 10.1371/journal.pone.0148441. PubMed PMID: 26863147; PubMed Central PMCID: PMCPMC4749302.

29. Schommer K, Kallenberg K, Lutz K, Bartsch P, Knauth M. Hemosiderin deposition in the brain as footprint of high-altitude cerebral edema. Neurology. 2013;81(20):1776–9. Epub 2013/10/11. doi: 10.1212/01.wnl.0000435563.84986.78. PubMed PMID: 24107867.

30. Kallenberg K, Dehnert C, Dorfler A, Schellinger PD, Bailey DM, Knauth M, et al. Microhemorrhages in nonfatal high-altitude cerebral edema. Journal of cerebral blood flow and metabolism : official journal of the International Society of Cerebral Blood Flow and Metabolism. 2008;28(9):1635–42. Epub 2008/06/05. doi: 10.1038/jcbfm.2008.55. PubMed PMID: 18523438.

31. Hempel C, Combes V, Hunt NH, Kurtzhals JA, Grau GE. CNS hypoxia is more pronounced in murine cerebral than noncerebral malaria and is reversed by erythropoietin. The American journal of pathology. 2011;179(4):1939–50. doi: 10.1016/j.ajpath.2011.06.027. PubMed PMID: 21854739; PubMed Central PMCID: PMCPMC3181359.

32. Penet MF, Viola A, Confort-Gouny S, Le Fur Y, Duhamel G, Kober F, et al. Imaging experimental cerebral malaria in vivo: significant role of ischemic brain edema. The Journal of neuroscience : the official journal of the Society for Neuroscience. 2005;25(32):7352–8. Epub 2005/08/12. doi: 10.1523/JNEUROSCI.1002-05.2005. PubMed PMID: 16093385.

33. Krueger M, Mages B, Hobusch C, Michalski D. Endothelial edema precedes blood-brain barrier breakdown in early time points after experimental focal cerebral ischemia. Acta Neuropathol Commun. 2019;7(1):17. Epub 2019/02/13. doi: 10.1186/s40478-019-0671-0. PubMed PMID: 30744693; PubMed Central PMCID: PMCPMC6369548.

34. Turner G. Cerebral malaria. Brain Pathol. 1997;7(1):569–82. PubMed PMID: 9034566.

35. McKinney AM, Sarikaya B, Gustafson C, Truwit CL. Detection of microhemorrhage in posterior reversible encephalopathy syndrome using susceptibility-weighted imaging. AJNR American journal of neuroradiology. 2012;33(5):896–903. doi: 10.3174/ajnr.A2886. PubMed PMID: 22241378.

36. Akoudad S, Wolters FJ, Viswanathan A, de Bruijn RF, van der Lugt A, Hofman A, et al. Association of Cerebral Microbleeds With Cognitive Decline and Dementia. JAMA Neurol. 2016;73(8):934–43. doi: 10.1001/jamaneurol.2016.1017. PubMed PMID: 27271785; PubMed Central PMCID: PMCPMC5966721.

37. Scheid R, Walther K, Guthke T, Preul C, von Cramon DY. Cognitive sequelae of diffuse axonal injury. Arch Neurol. 2006;63(3):418–24. doi: 10.1001/archneur.63.3.418. PubMed PMID: 16533969.

38. Valenti R, Del Bene A, Poggesi A, Ginestroni A, Salvadori E, Pracucci G, et al. Cerebral microbleeds in patients with mild cognitive impairment and small vessel disease: The Vascular Mild Cognitive Impairment (VMCI)-Tuscany study. J Neurol Sci. 2016;368:195–202. doi: 10.1016/j.jns.2016.07.018. PubMed PMID: 27538632.

39. Carroll RW, Wainwright MS, Kim KY, Kidambi T, Gomez ND, Taylor T, et al. A rapid murine coma and behavior scale for quantitative assessment of murine cerebral malaria. PloS one. 2010;5(10). Epub 2010/10/20. doi: 10.1371/journal.pone.0013124. PubMed PMID: 20957049; PubMed Central PMCID: PMC2948515.

40. Mastronarde DN. Automated electron microscope tomography using robust prediction of specimen movements. J Struct Biol. 2005;152(1):36–51. doi: 10.1016/j.jsb.2005.07.007. PubMed PMID: 16182563.

41. Ng L, Bernard A, Lau C, Overly CC, Dong HW, Kuan C, et al. An anatomic gene expression atlas of the adult mouse brain. Nature neuroscience. 2009;12(3):356–62. Epub 2009/02/17. doi: 10.1038/nn.2281. PubMed PMID: 19219037.

42. Schneider CA, Rasband WS, Eliceiri KW. NIH Image to ImageJ: 25 years of image analysis. Nature methods. 2012;9(7):671–5. Epub 2012/08/30. PubMed PMID: 22930834.

